# Aldehyde dehydrogenase protein: a placental glycogen trophoblast marker suitable to monitor placental pathologies

**DOI:** 10.1101/2025.09.01.673566

**Authors:** Alka Rani, Xinrui Li, Melissa Arboleda, Heidi Stuhlmann

## Abstract

This study focuses on developing new approaches to identify and quantify glycogen trophoblast cells in the placenta. Due to limitations with currently available methods, alternative markers for these cells are being explored. Here we report on one promising candidate, *Aldh1a3*, a gene that was recently shown to be expressed in the mouse placenta only in glycogen trophoblast cells and their progenitors. Our study validates ALDH1A3 protein as glycogen trophoblast marker in the mouse placenta with high specificity when compared to CDKN1C (p57KIP2). This marker will be useful for the isolation of the glycogen trophoblast subpopulation and facilitate examining its function in placental programming and understanding its role in pathological pregnancy models. Furthermore, trophoblast giant cells with high expression of ALDH1A3 were detected in the decidua of human term placentas. These were interspersed with extravillous interstitial trophoblast cells or were located distal of the interstitial trophoblast layer of the decidua basalis.

## Introduction

The mouse placenta has a unique structure compared to the human placenta, making it important to have unique markers to differentiate between different trophoblasts. Glycogen trophoblast is one such unique trophoblast subtype that has specific roles during the mouse placental development. It is not only important as a storage system for glycogen to support the nourishment for development of the embryo, but also possesses endocrine functions. It is located in the junctional zone which forms a barrier between the uterine wall and the placental labyrinth. Glycogen trophoblast cells have been shown to be affected in pathological pregnancy, gene knockout, and metabolic insult models in mice. Therefore, identifying glycogen trophoblast-specific biomarkers that can be used with 100% specificity is an important task.

The few markers that have been identified for glycogen trophoblasts in the mouse placenta include CDKN1C (p57KIP2) and different members of the Prolactin gene family. Previously, we reported the expression pattern of CDKN1C protein in the mouse placenta (1). Since CDKN1C is located in the nucleus, its use to identify glycogen trophoblast in the junctional zone requires co-staining with placental structural markers like trophoblast-specific protein alpha (TPBPA). Therefore, we were interested to explore and validate novel glycogen trophoblast specific markers that can be used independently.

Recently, Aldh1a3 mRNA was shown to be exclusively expressed in glycogen trophoblast cells in the mouse placenta and their progenitors in the ectoplacental cone, using in-situ hybridization (2, 3). ALDH1A3 is an enzyme involved in the biosynthesis of retinoic acid, which plays crucial roles in various developmental processes. To validate its potential as a glycogen trophoblast-specific marker, further studies are needed to assess its protein expression patterns and develop a robust method convenient for researchers.

Our study aimed to provide a comprehensive analysis of ALDH1A3 protein localization and to compare its localization with that of CDKN1C in various murine placental regions. Additionally, we used our established *miR-126* knockout mouse model that displays junctional zone hyperplasia due to increased numbers of proliferating glycogen trophoblast cells (1) to further validate that ALDH1A3 can serve as a biomarker to study glycogen trophoblasts pathology in the placenta. Furthermore, we localized ALDH1A3 in the human term placenta in the basal plate.

## Materials and Methods

### Placenta Collection

All animal protocols were approved by the Institutional Animal Care and Use Committee (IACUC) at Weill Cornell Medical College of Cornell University. Timed pregnancies of C57BL/6 or miR-126 females crossed with male mice (morning of vaginal plug was counted as E0.5) were set up, and placenta samples (n=5 each) were collected at gestational day E15.5 and E18.5, respectively. Placentas were fixed in 4% paraformaldehyde (PFA) for 2–4 h at 4°C with gentle agitation, embedded in OCT and stored at −80°C until further analysis.

Human placenta samples were collected from healthy term pregnancies (n=3) shortly after delivery under an institutional IRB-approved protocol. The samples were fixed overnight at 4°C in 4% paraformaldehyde (PFA), washed the next day in PBS, placed overnight at 4°C in 30% sucrose/PBS, and embedded in a 1:2 mixture of 30% sucrose in OCT. All the samples were stored at −80°C until analysis.

### Immunofluorescence

For sectioning, blocks were equilibrated at −20°C and cryo-sectioned at 8-10 µm thickness. Sections were mounted on Superfrost™ Plus slides and air-dried. Slides were equilibrated to room temperature (RT) for 1 h, post-fixed in cold acetone (−20°C) for 10 min, and dried at 60°C for 5 min. Antigen retrieval was performed by heating in sodium citrate buffer (pH 6.0) at 98–100°C for 5 min. Slides were blocked with 5% donkey and 5% goat serum in PBST (PBS + 0.1% Tween-20) for 1 h, followed by overnight incubation at 4°C with first primary antibodies diluted in PBST. After PBST washes, fluorescently labeled secondary antibodies were applied for 1 h at RT in the dark. After washing the slides were incubated with second primary antibody for 1hr at RT, followed by second secondary antibody incubation as described before. ProLong Gold Antifade Mountant with DNA Stain DAPI (Invitrogen: P36931) was added and cover slips were placed. Slides were cured overnight at 4°C and imaged the next day.

Antibodies used included CD31 (1:100, R&D: AF3628, 0.2 mg/mL), ALDH1A3 (1:200, Novus Biologicals: NBP2-15339, 1.0 mg/ml), p57KIP2 (1:200, CDKN1C) (Proteintech: 23317-1-AP, 433 ug/ml), and KRT7 (1:500, Agilent Technologies, M701829-2). The ALDH1A3 and CDKN1C antibodies were from same rabbit host species; therefore, AffiniPure Fab Fragment Goat Anti-Rabbit IgG (H+L) (Jackson Lab. Inc.:111-007-003, 1.3 mg/ml) was used between the two antibody incubations to avoid cross linking of the secondary antibodies (4). Secondary antibodies used for mouse placenta staining included Alexa488-donkey-α-rabbit and Alexa594-goat-α-rabbit (Jackson Immunoresearch, 1.5□µg/ml) at 1:400 dilution. Prior to IF staining of human placentas, antigen retrieval was performed with sodium citrate buffer (pH 6.0) at 98-100°C for 5 min, followed by blocking with 5% donkey and 5% goat serum in PBST (PBS + 0.1% Tween-20) and incubation with primary antibodies. Alexa Fluor 488 donkey anti-rabbit IgG (ThermoFisher, 2 mg/mL; 1:1000 dilution) and Alexa Fluor 594 goat anti-mouse IgG (Jackson ImmunoResearch, 1.5 µg/mL; 1:400 dilution) were used as secondary antibodies. Images were acquired and whole mouse placenta was scanned using a Zeiss Axiovert automated fluorescence microscope and ZEN software at 10X and 20X. Images of human placenta sections were acquired using a Zeiss AxioImager Z1 fluorescence microscope equipped with an AxioCam MRm camera at 10X and 20X and AxioVision software version 4.8. Exposure times were calibrated using negative controls to minimize background fluorescence and ensure consistent image quality.

## Results and Discussion

To locate expression of ALDH1A3 protein in glycogen trophoblast cells of the mouse placenta, we co-stained sections from wild-type E18.5 placentas with antibodies for ALDH1A3 and the previously described marker CDKN1C, a nuclear marker expressed in glycogen trophoblast but not in spongiotrophoblast cells of the junctional zone (1, 5, 6). ALDH1A3 and CDKN1C staining co-localized to the same cells in clustered areas of placental glycogen trophoblasts in the junctional zone and the decidua (Fig. 1A-C). ALDH1A3 was localized in the cytoplasm, whereas CDKN1C was more diffusely located in the nucleus. ALDH1A3 expression was not detected in the labyrinth zone (LB), whereas expression of CDKN1C was observed in a subset of labyrinth cells consistent with its previous reported expression in syncytiotrophoblast (7) (Fig. 1D).

**Figure 1:**
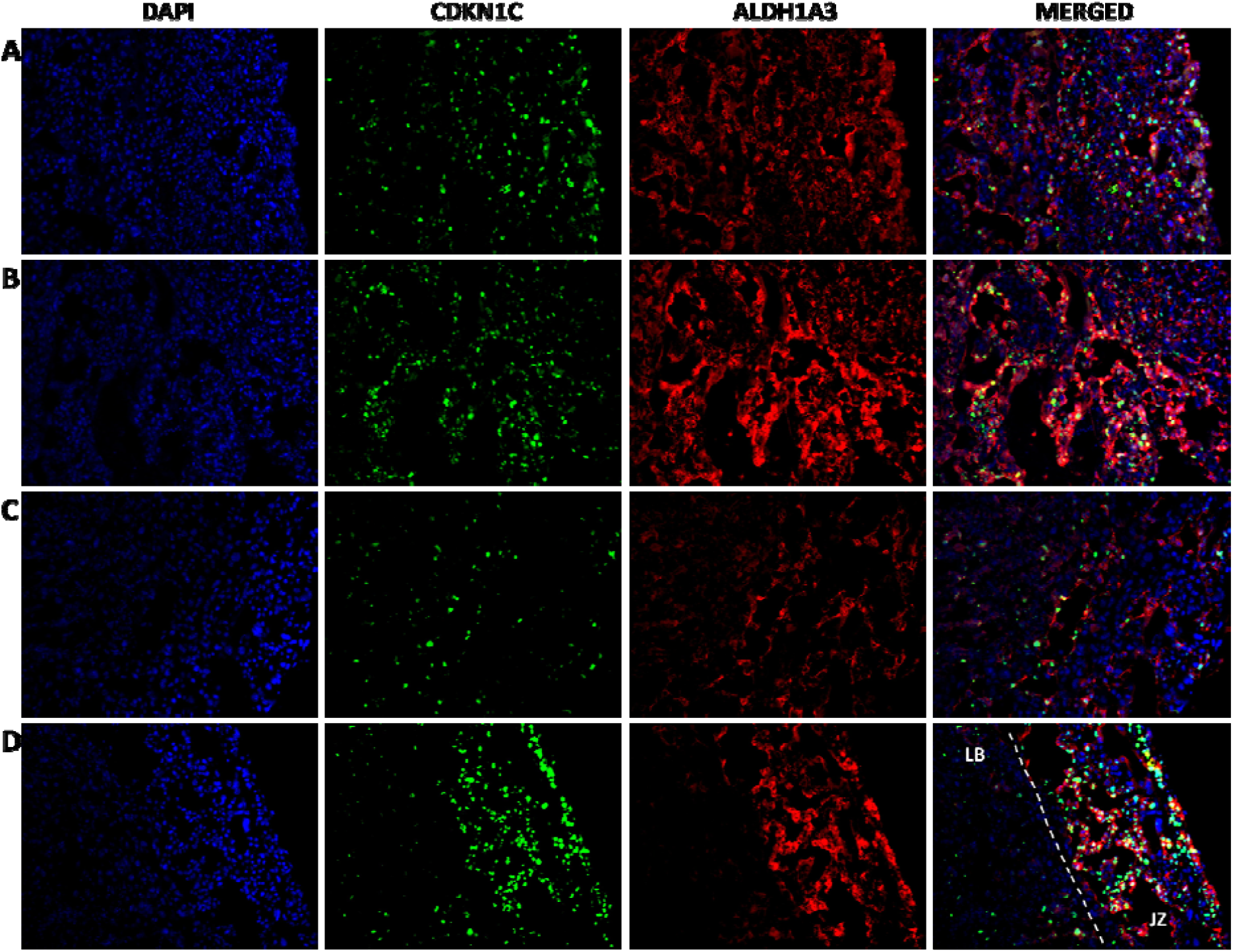
Co-localization of CDKNI1C and ALDH1A3 Proteins in the Mouse Placenta. Immunostaining showing co-localization of CDKN1C (green) and ALDH1A3 (red) protein in glycogen trophoblasts in the junctional zone region of the E18.5 mouse placenta. (A, B) Strongly stained regions (20X); (C) less stained region at 20X; (D) CDKN1C and ALDH1A3 co-stain in the junctional zone (JZ); ALDH1A3 staining is restricted to the JZ of the placenta, in contrast to CDKN1C (10X).

Glycogen trophoblasts are integral to placental function and fetal development in mice, particularly under pathological conditions. Glycogen trophoblast cells in the mouse placenta emerge from the spongiotrophoblast layer and accumulate glycogen in storage vesicles. Around E14.5, a subpopulation of glycogen trophoblast cells migrates into the maternal decidua (8). Highest numbers of glycogen trophoblast cells are detected at E15.5, and between E16.5 and E18.5 the number of glycogen trophoblasts starts reducing when they migrate closer to the labyrinth and junctional zone barrier of the placenta, as shown in Fig. 1D. To visualize the localization of ALDH1A3-positive glycogen trophoblast cells in E15.5 placentas, we performed co-staining using CD31 as a marker for fetal endothelial cells that are restricted to the labyrinth region (Fig. 2). ALDH1A3 protein was observed in the junctional zone and decidua regions of the 15.5 mouse placenta but was absent in the CD31-positive area of the fetal labyrinth, establishing that ALDH1A3 marks specifically glycogen trophoblast cells in the murine placenta. Our results further indicate that ALDH1A3, as a cytoplasmic marker, provides a better visual understanding of the clustering and the spreading of glycogen trophoblast cells between E15.5 and E18.5 in the mouse placenta.

**Figure 2:**
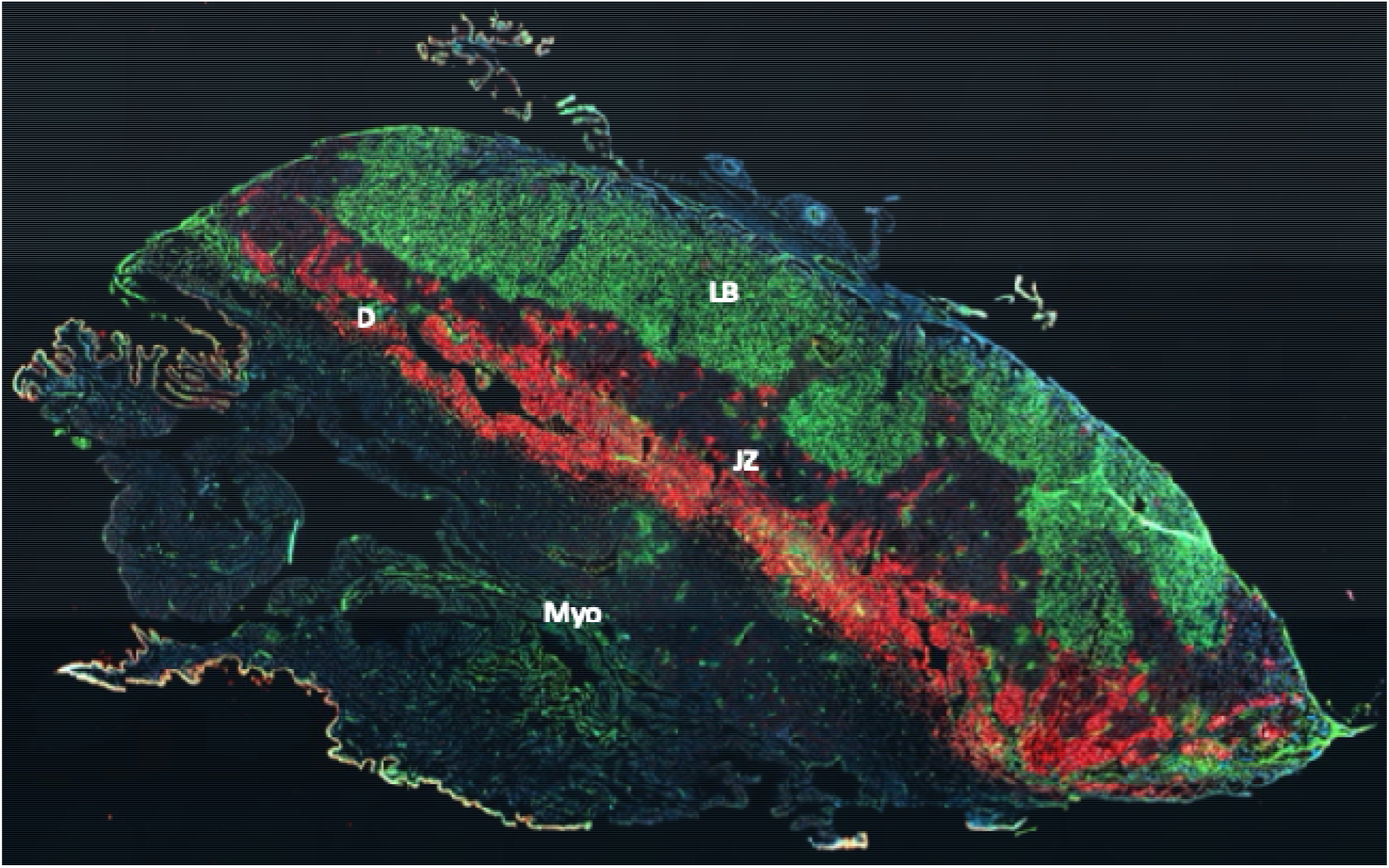
ALDH1A3 staining in E15.5 Mouse Placenta. Immunostaining showing CD31 (green) protein marking the labyrinth region (LB) and ALDH1A3 (red) staining the glycogen trophoblast clusters spreading from the junctional zone (JZ) into the decidua (D) of the E15.5 mouse placenta.

Furthermore, co-staining using ALDH1A3 together with CDKN1C antibodies, with the help of using the Fab fragment as they are both raised in the rabbit as the host species, can also help researchers for quantification analysis by counting CDKN1C-positive nuclei overlayed in the ALDH1A3 stained regions of the placenta. Quantification of CDKN1C-positive nuclei requires an overlay of junctional zone marker, such as TPBPA, to be able to identify the nuclei of glycogen trophoblasts (1). As shown in Fig. 3A, CDKN1C also stains non-glycogen trophoblast nuclei in the labyrinth region of the placenta. Of note, when TPBPA antibody supply stopped over the past few years in United States, we tested several antibodies specific for glycogen trophoblast and junctional zone cells, including Connexin 31 (GJB3) and Prolactin, but none of these provided satisfactory results in our lab (not shown). When we used ALDH1A3 antibody the problem was solved, as it stained glycogen trophoblasts of high specificity and with no staining in the labyrinth region of the placenta as compared to CDKN1C (Fig. 3A&B). Importantly, *mir-126* knockout placentas were used here for validation, as they contain increased size of glycogen trophoblast cell clusters increased number of glycogen trophoblasts in the junctional zone (1). Results consistent with those previous studies are presented here by staining for both CDKN1C and ALDH1A3 markers (Fig. 3). In summary, we show that ALDH1A3 is a highly efficient and specific marker for glycogen trophoblast cells that it can be used in combination with CDKN1C for quantitative studies.

**Figure 3:**
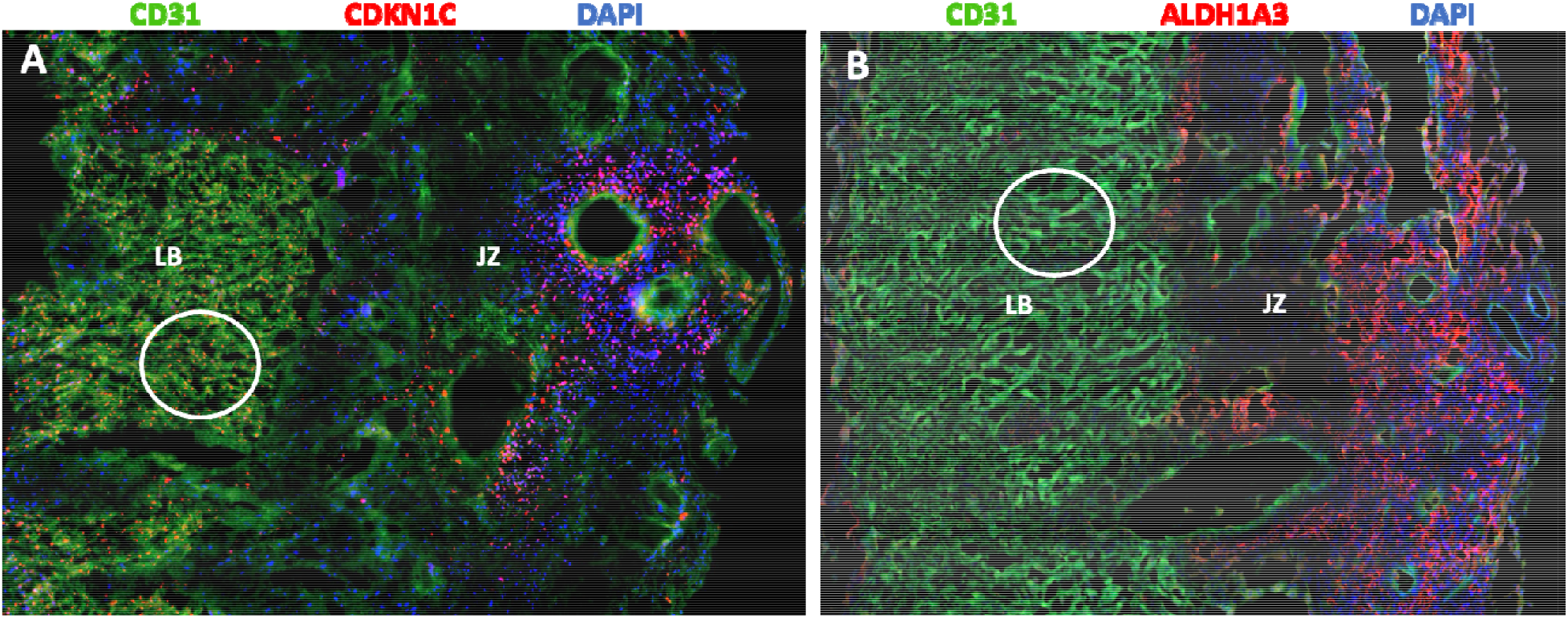
CDKN1C and ALDH1A3 Staining in Labyrinth Region in the Pathological Placenta. Immunostaining showing CDKN1C (red) staining in glycogen trophoblast cells of the junctional zone (JZ) and of trophoblast cells of the labyrinth region (LB) that is marked by CD31 (green) (red dots inside the white circle) in E15.5 *mir-126*^−/−^ mouse pathological placenta. (B) showing absence of ALDH1A3-positive cells (red) in the LB region (no red stain inside the white circle). Junctional zone (JZ) containing glycogen trophoblasts show expression of both CDKN1C (A) and ALDH1A3 (B) markers.

The role of glycogen trophoblast extends beyond energy storage; they are endocrine cells that are implicated to play roles in glucose release and placental remodeling, processes essential for fetal growth. Alterations in glycogen trophoblast dynamics have been observed in various mouse models of pregnancy complications. For instance, in *miR-126* knockout mice, we observed increase in glycogen trophoblast numbers correlates with intrauterine growth restriction (IUGR) and reduced fetal viability, highlighting the impact of glycogen trophoblast proliferation on placental architecture and function (1). Additionally, factors like insulin-like growth factor 2 (IGF2) and granulocyte–macrophage colony-stimulating factor (CSF2) influence glycogen trophoblast development and function, suggesting that disruptions in these pathways can lead to placental insufficiencies (9). Therefore, understanding the behavior and regulation of glycogen trophoblast is crucial for elucidating the mechanisms underlying placental pathologies and fetal development in mouse models.

To identify the corresponding glycogen trophoblast cells in the human placenta, we performed immunofluorescence on sections from healthy human placenta samples from term pregnancies. While the human placenta does not contain specialized glycogen trophoblast cells like mice, glycogen is stored in the decidual cells of the uterine lining (decidua) and is crucial for supporting the early stages of implantation and trophoblast invasion. The glycogen-rich decidua provide an optimal environment for early placental growth by supplying energy (10). During early pregnancy, extravillous cytotrophoblast cells are organized in columns and are located at the basal plate of the placenta. Invasion of the decidua basalis by cytotrophoblast-derived interstitial trophoblast cells is underway as early as 7 weeks of gestation, and subsequent invasion of inner layer of the myometrium is completed by 22 weeks (6). Studies on human placentas from different gestational stages suggested that vacuolized, glycogen-containing trophoblast cells in the decidua basalis and myometrium are derived from invading interstitial trophoblast cells. Co-staining of sections from three healthy term placentas with ALDH1A3 and KRT7, a well-characterized marker for trophoblast cells, provided a dynamic picture of the emergence of glycogen metabolizing cells in the decidua basalis (Fig. 4A-C). Individual cells and small clusters of invading interstitial trophoblasts are visible in the decidua basalis as small, sphere-shaped KRT7^+^ ALDH1A3^−^ cells and are interspersed with decidual cells KRT7^−^ ALDH1A3^+^ (Fig. 4A-C). In some images, the KRT7^+^ trophoblasts have changed their shape to become giant trophoblast cells co-expressing ALDH1A3 (Fig. 4B), suggesting that these undergo differentiation into glycogen trophoblast cells. Finally, a few chorionic villi showed weak expression of ALDH1A3 in syncytiotrophoblast cells (Fig. 4B).

**Figure 4:**
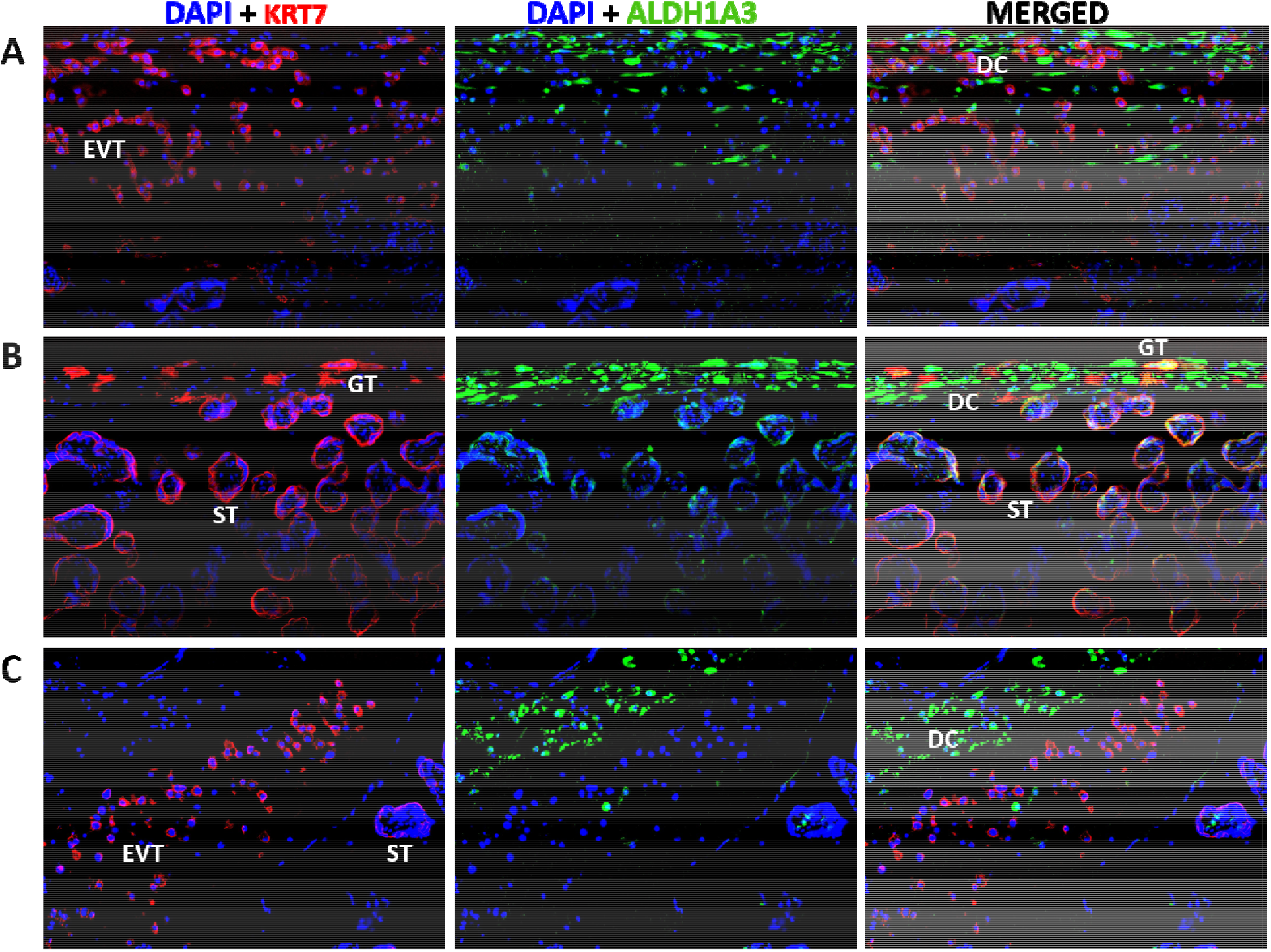
ALDH1A3 Proteins Localization in Human Term Placenta. Immunostaining of human term placentae basal plate showing ALDH1A3 (green) stained decidual cells (DC) and extra villous trophoblast (EVT) only stained for trophoblasts marker cytokeratin 7 (KRT7; red) (A-C). Giant trophoblast cells (GT) and syncytiotrophoblast (ST) stained for both ALDH1A3 (green) and KRT7 (red) shown in the merged column (yellow) (B). 20X magnification images.

ALDH1A3 expression was consistently strong in the decidual cells. The histology of decidual cells and extravillous trophoblasts have been well characterized in the literature, as shown in the illustration Fig. 5A (11). Therefore, we performed H&E staining on sections of human placentas adjacent to those shown in Fig. 4. Comparing the staining patterns confirmed the identity of KRT7^−^ ALDH1A3^+^ decidual cells with spindle to oval shape containing elongated nuclei that are forming the top layer of the basal plate. Clusters of KRT7^+^ ALDH1A3^−^ cells were confirmed to be extravillous trophoblasts with similar large polygonal cells and large round nuclei found below and embedded in the decidual cells of the basal plate (Fig. 5B). Future studies will be required to explore the role ALDH1A3 plays in the decidual cells of the human placenta at term.

**Figure 5:**
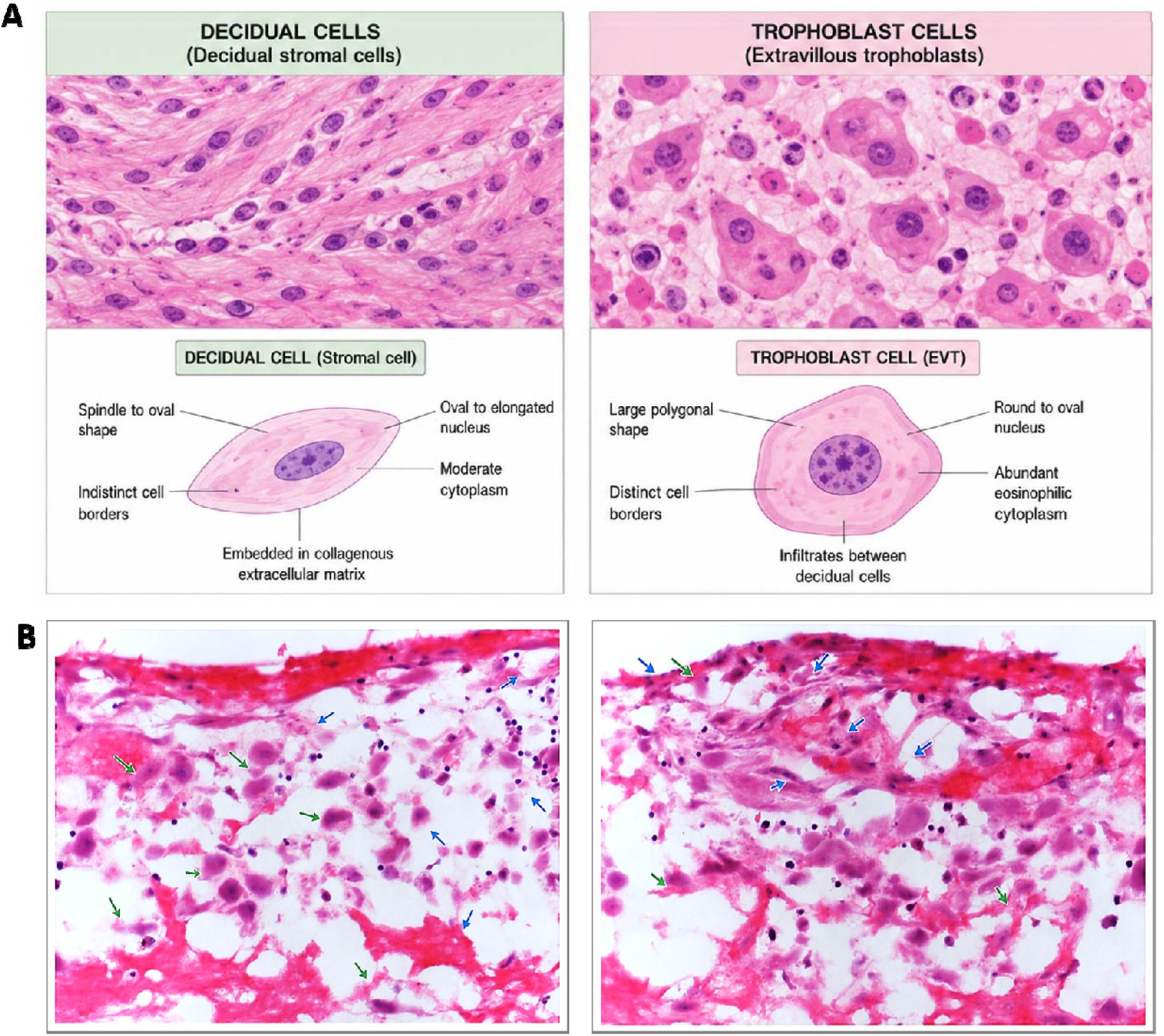
Distribution of Decidual cells and Trophoblast cells in Human Term Placenta. AI generated illustration showing characteristics of decidual cells (DC) and extravillous trophoblasts (EVT) identified in the human placenta (A). H&E staining of sections adjacent to those in Fig. 4, showing distribution of spindle to oval cells with elongated nuclei DC (Blue Arrow) and large polygonal cells with abundant cytoplasm EVT (Green Arrow) in the basal plate of term human placentae (B). 20X magnification images.

In conclusion, we validated in this study the presence and restricted localization of ALDH1A3 protein in mice and human placentae for the first time. ALDH1A3 was observed to be a highly specific marker for glycogen trophoblasts, making it a highly useful tool to study placental pathologies in mouse models.

